# A Phylogenetic Perspective on the Syntax-Semantics Interface: A Case Study in Clause Linkage Constructions

**DOI:** 10.1101/2025.07.19.665697

**Authors:** Carlo Y. Meloni, Chundra A. Cathcart, Erik J. Ringen, Jessica K. Ivani, Balthasar Bickel

**Affiliations:** Institute for the Interdisciplinary Study of Language Evolution (ISLE), University of Zurich; Linguistic Research Infrastructure (LiRI), University of Zurich; Department of Linguistics, University of Tübingen; Department of Linguistics, University of California, Santa Barbara

**Author notes:** These authors contributed equally to this work (Jessica K. Ivani, Balthasar Bickel).

## Abstract

We examine the syntax-semantics interface in clause linkage using a cross-linguistic sample of nearly 700 constructions from Indo-European, Sino-Tibetan, and Tupí-Guaraní. To evaluate predictions from the Interclausal Relations Hierarchy, we model the phylogenetic dynamics of how semantic relations influence the degree of syntactic integration between clauses. The best performing model assumes an Ornstein-Uhlenbeck process in each family and suggests only moderate and lineage-specific effects of semantics on syntax, challenging the universality of the Interclausal Relations Hierarchy. Furthermore, geographic proximity exerts only minimal effects, consistent with the notion that the syntax-semantics interface may be more resistant to contact effects than overtly signaled and transparent patterns in language. Together, our findings suggest that the syntax-semantics interface is less constrained in clause linkage than is commonly assumed in typology. They furthermore highlight the value of treating phylogeny as a dynamic evolutionary process, rather than as a static background structure.

## 1 Introduction

The strategies that languages employ in combining event-denoting utterances, a phenomenon known as clause linkage, exhibit extreme cross-linguistic and within-language variation. For example, many languages use the same construction for both direct and indirect speech, whereas others require the overt use of a complementizer when reporting indirect speech. Some languages mark illocutionary scope over both clauses, whereas similar-looking constructions in other languages assume the truth value of one of the clausal propositions [1, 2]. Despite this variation, a number of cross-linguistic generalizations have been proposed regarding the nature of clause linkage, particularly with respect to the relationship between the semantic properties of a construction and the syntactic structure it exhibits. Specifically, many linguists have proposed that there is a correlation between the syntactic and semantic integration of clause linkage structures. Yet this hypothesis has not been investigated on a wide scale using state-of-the-art quantitative methods.

In this paper, we assess the evidence for an association between semantic closeness and syntactic integration, basing our analyses off of clause linkage hierarchies proposed in the literature. We use fine-grained data comprising information about asymmetries between clauses in a large number of constructions from languages spanning three families. We analyze these constructions using statistical models capable of detecting associations between variables while accounting for phylogenetic and spatial non-independence, as well as statistical models that explicitly model dynamics of change in each phylogeny. Our best fitting models support a moderate relationship between syntactic and semantic integration, although the detailed nature of this relationship strongly differs across families. We conclude with a discussion of the methodological implications of our findings for future work concerning statistical modeling in typological research.

## 2 Background

### 2.1 Theoretical proposals on the syntax-semantics interface

Clause linkage has long been a central topic in linguistic research. Over the past few decades, the typological literature has increasingly focused on the relationship between the structural and semantic properties of clause-combining constructions. One area of particular interest is the degree of integration between clauses: how tightly clauses are bound together syntactically and semantically, and how this relates to notions such as clause union and event boundedness.

Syntactic integration refers to the structural closeness of combined clauses. For example, in a sentence like ‘I want to finish the paper’, the complement clause ‘to finish the paper’ is syntactically integrated with the main clause in several ways: the main clause subject controls the zero subject of the dependent clause; the embedded verb is non-finite (‘to finish’), and the sentence typically forms a single intonational unit [3]. Semantic integration, on the other hand, pertains to the conceptual or event-level relationship between clauses. In the same example, event boundedness is reflected by subject co-reference and by the main verb (‘want’) expressing the subject’s attitude (desire and intent) toward the embedded event (finishing the paper).

Numerous accounts have argued for a systematic correlation between syntactic and semantic integration. According to this view, the closer two events are in meaning and cognitive representation, the more tightly they tend to be linked syntactically. This correlation is often interpreted as a form of iconicity, where the formal properties of a construction mirror the nature of the conceptual relationship it encodes. In what follows, we briefly review key contributions to this proposal.^1^

One of the earliest and strongest proponents of this view is Givón [13, 14, 3], who explains syntactic patterns in clause combining, especially complementation, in terms of systematic syntactic-semantic isomorphism. In addition to identifying individual prototypes of syntactic and semantic isomorphisms, Givón argues that both dimensions are gradual and scalar across complement clause types, reflecting diachronic processes. On one end are perception/cognition and speech utterances, characterized by looser semantic relations and typically realized as structurally independent clauses; on the other end are conceptually dependent events (such as those in causative, manipulative, or modal complement constructions) which tend to be syntactically compressed.^2^ These distributions are explained by invoking functional phenomena, such as iconicity.

Cristofaro [19] broadens the cross-linguistic scope to all types of subordination, including adverbial, complementation, and relative relations. These constructions are formally described according to cross-linguistic comparable parameters and are semantically categorized based on the cognitive and functional relation between events in the construction. The main empirical result is the *Subordination Deranking Hierarchy*, which predicts, among other things, that semantic relations such as phasal, modal, and manipulatives tend to be encoded using more syntactically integrated forms, compared to other relations (for example, those capturing knowledge or propositional attitudes).

Van Valin [20], building on earlier work, such as Foley and Van Valin [1], provides perhaps the most comprehensive theoretical treatment of the syntax-semantics interface. The *Interclausal Relations Hierarchy* (IRH) explicitly links semantic relations between clauses with syntactic configurations. Semantic relation types are arranged on a hierarchical continuum of semantic cohesion (in the *Interclausal Semantic Relations Hierarchy*), ranging from loosely connected propositional units (e.g., successive temporal clauses, characterized by multiple events) to tightly cohesive relations (e.g., phase and manner, representing facets of a single event). Syntactic configurations, parametrized along structural properties, are similarly ranked by the strength of their syntactic bond. The IRH aligns these two hierarchies based on the principle that the closer the semantic relationship between two propositions, the stronger the syntactic link joining them. Van Valin embeds this correlation within the formal architecture of grammar through a mechanism of linking analogues to how semantic roles are linked to syntactic functions. The IRH maps semantic representations in event composition to syntactic structure in clause linkage expression.

Taken together, these works suggest that clause linkage reflects systematic relationships between syntactic structure and semantic meaning, whether explained through iconicity or formal linking mechanisms in grammar. Yet, most studies to date have been qualitative, and robust cross-linguistic analyses require systematic ways to represent these features in data. In parallel with these theoretical proposals, there has been an increasing effort to formalize and parametrize the structural and semantic properties of clause combining for cross-linguistic comparison. In what follows, we introduce major approaches to the parameterization and data structuring of clause linkage in the literature.

### 2.2 Data structuring in clause linkage

The IRH structures syntactic configurations around two key parameters. The first is nexus relations, which describe the syntactic relations between the units in a complex construction. This parameter allowed for the introduction of cosubordination [21, 1, 20] as a third nexus type alongside the traditional concepts of subordination and coordination. The second key parameter concerns the nature of the units being linked (referred to as juncture in the Role and Reference Grammar framework), distinguishing between core, nuclear, and clausal juncture types.

Based on the interaction of these parameters, Van Valin [22] identifies eleven abstract juncture-nexus types,^3^ which can be arranged hierarchically according to the strength of the syntactic bond: from nuclear cosubordination (strongest) to sentential coordination (weakest). The eleven juncture-nexus types, while purely syntactic, are used to express certain semantic relations between the units in the juncture.

Another important contribution to the structural parameterization of clause combining is found in Lehmann [23]. His typology identifies clause linkage types based on the interaction of several semanticosyntactic parameters, including desententialization, main syntactic level, hierarchical downgrading of the subordinate clause, grammaticalization properties of the main verb, interlacing, and the explicitness of the linking. Each parameter (except interlacing) is organized along a continuum, broadly paralleling the nexus classification described above, but with some crucial differences in descriptive granularity. In Van Valin [22], the eleven juncture-nexus types refer to abstract clause-combining types taken as a bound whole (e.g., ‘core subordination’); Lehmann’s framework, by contrast, introduces finer distinctions by de-structuring the syntactic domain. For example, while parameters like hierarchical downgrading use whole clause types in their continuum (such as ‘adjoined’ or ‘medial clauses’), other variables, like the extent of grammaticalization of the main verb, refer to more granular properties within the construction.

The typological generalizations in Cristofaro [19], including the *Subordination Deranking Hierarchy*, are grounded in cross-linguistically viable formal and semantic/functional descriptions. Two methodological advances are central to this approach: first, key properties of subordination constructions (such as finiteness of the subordinate clause) are defined in terms of the language-specific structural distance from the respective independent declarative clause; second, functional criteria, especially types of semantic relations, are placed at the forefront. Subordination is thus characterized in functional terms, and subordination strategies are categorized according to the cognitive and functional relationship between events (state of affairs) in the construction.

Building on the need for greater granularity in describing clause-linking constructions, Bickel [2] proposes a multivariate typological approach that analyzes each construction as a set of independent structural variables. This approach allows both language-specific descriptions and systematic cross-linguistic analysis and moves away from predefined types. One empirical outcome is a pilot database of 69 constructions from 24 languages, coded for 11 structural variables capturing both syntactic and semantic properties of combined clauses. These variables include the scope of operators (tense, negation, and illocutionary scope), degree of finiteness, symmetry between dependent and main clause (also emphasized in Haiman, Haiman, and is connected to the distinction between balanced vs. deranked in Cristofaro [19], position of the dependent clause, degree of embedding (layering), focus marking, and extraction possibilities from the dependent clause.

This construction-based approach is further developed in van Gijn, Galucio, and Nogueira [24]’s typological study of subordination strategies in Tupian languages. Their analysis is grounded in the principles of Construction Grammar [25], treating clause-linkage patterns as constructions, that is, form-meaning pairings analyzed along both formal and semantic dimensions. Building on Cristofaro [19], van Gijn, Galucio, and Nogueira [24] examine subordination strategies primarily from a semantic perspective, with the units of description defined by how languages encode specific semantic relations between state of affairs (or *event-denoting units*, henceforth EDU). This broader approach encompasses not only traditional subordinating constructions but also nominalizations, serial verb constructions, auxiliary verb constructions, and verb-verb compounds expressing similar relations. The formal properties of each construction are described across five key morphosyntactic domains (detailed in Section 3), and are inspired by previous studies, especially Lehmann [23], Cristofaro [19], and Bickel [2]. A significant component of the questionnaire by van Gijn, Galucio, and Nogueira [24] involves annotating the semantic relations expressed by clause linkage constructions. These semantic subordination relations, drawn on Cristofaro [19], include several functions, including temporal (simultaneous, successive), causal, locational, phasal, modal, perceptual and reportative, among others. Each semantic relation is coded as an independent variable, which captures the versatility or specialization of individual constructions.

## 3 Data

In our study, we describe clause linkage patterns through a construction-based approach, where the units of description are individual clause-combining structures. These are coded in a questionnaire following a multicategorization design principle [26], where each variable is coded through binary values.The structure of the questionnaire and the coding scheme follow van Gijn, Galucio, and Nogueira [24], but we adjusted the sets of variables during data transformation to ensure variable independence and better fit our research goals. The questionnaire captures five main morphosyntactic domains, outlined in van Gijn, Galucio, and Nogueira [24] and briefly introduced here.

One of the main domains is finiteness. Finiteness variables assess whether core verbal categories (like agreement, tense, aspect, and modality, among others) can be independently marked on the dependent clause, reflecting its degree of deverbalization. For example, the Albanian sentence in (1) explicitly marks subject agreement in the second EDU, whereas in the Kamaiurá gerund construction in (2), subject agreement appears only on the first predicate.

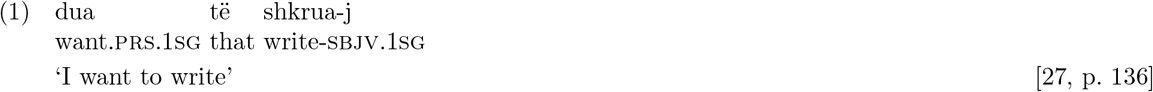

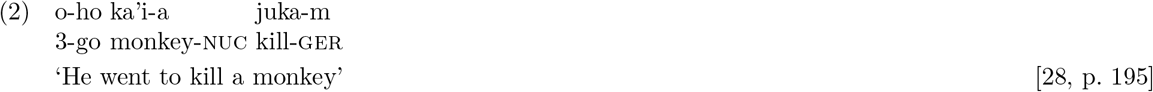

Nominalization variables describe the ‘noun-like’ properties of the dependent EDU, such as its ability to take case markers, determiners, or express possessors, among other things. An example from our data comes again from Kamaiurá, where the nominalized predicate in sentence (3) is marked with the nuclear case.

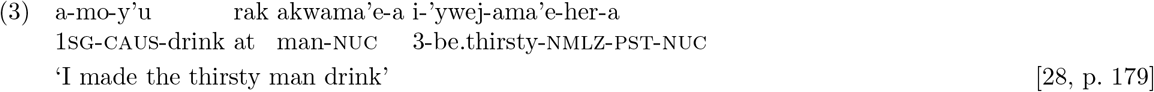

Dependency marking (or flagging) variables describe whether the dependent EDU is explicitly marked as subordinate, and where such marking occurs.

A section of the questionnaire is dedicated to relativization strategies, addressing how nominal modification is achieved with embedded clauses, including the marking and position of the relativized element. Sentences (4) and (5) illustrate variation in the linear ordering of relative clauses, with Mandarin placing the relative clause before the head noun, and Old High German placing it after.

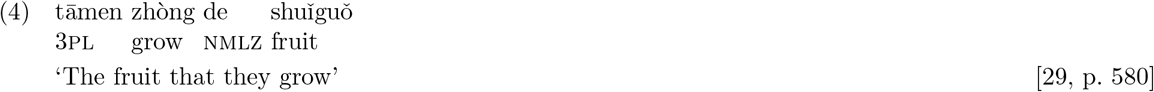

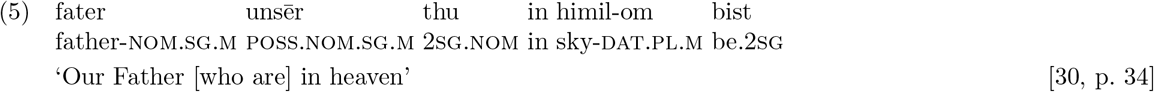

Another important set of parameters addresses the structural integration of clause combining. This includes items such as whether both EDUs can be negated independently. For instance, in Kokama juxtaposition (6), each EDU may carry negation markers, whereas in Lhasa Tibetan serial verb constructions (7), typically only one verb can be negated. Structural integration variables also describe whether the EDU must appear contiguous, and whether they are morphologically fused. These properties help determine how tightly bound the two EDUs are, ranging from loosely connected biclausal structures to tightly integrated compounds or affixed constructions.

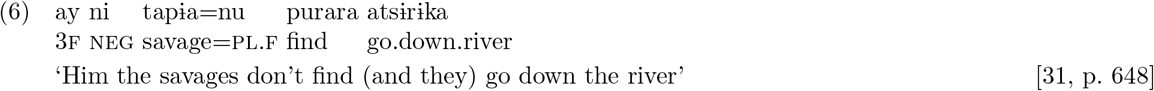

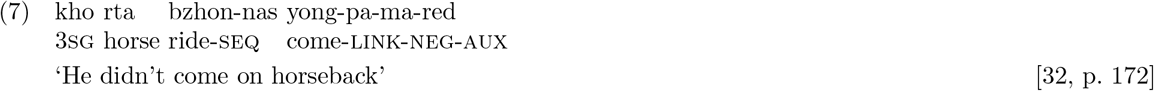

An additional parameter strictly related to this group is linearization, which captures word order preferences. Specifically, the parameter asks whether the dependent EDU precedes or follows the main clause, or whether the order between them is free.

The final part of the questionnaire describes the semantic relations that clause linkage constructions can encode. The semantic types relations included in the questionnaire are those adopted by van Gijn, Galucio, and Nogueira [33] and are drawn on an inventory of semantic types developed by Cristofaro [19]. These include variables assessing whether a construction can express temporal (simultaneous, successive), causal, locational, purposive, conditional (realis, counterfactual), phasal, modal, desiderative, manipulative, cognitive, perceptual, and reportative relations, among others. For instance, the Mandarin Chinese examples in (8) and (9) illustrate two different types of semantic relations: the causative construction in (8) expresses a causal meaning, whereas the relative clause in (9) encodes a locational relation.

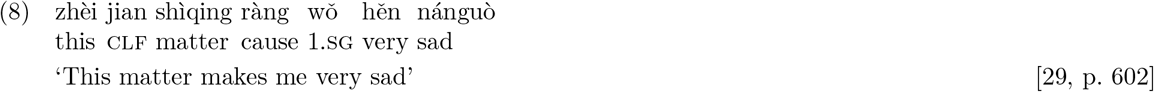

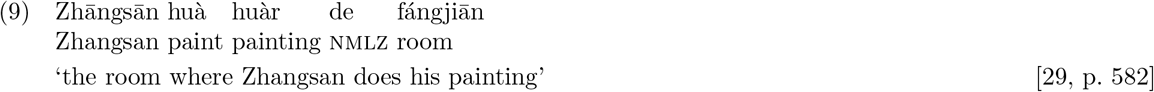

Each semantic function is treated as an independent variable, strengthening descriptive power. For example, semantic relation types may overlap, as shown in the following French examples:

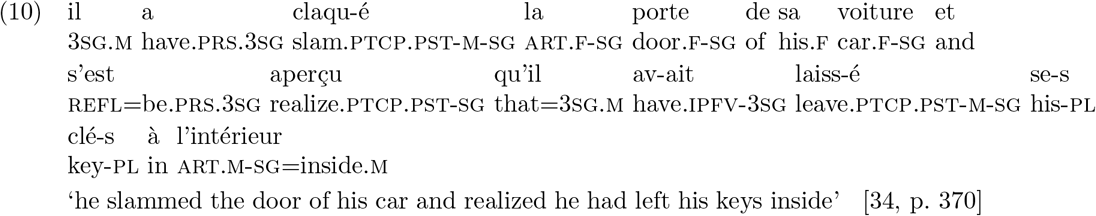

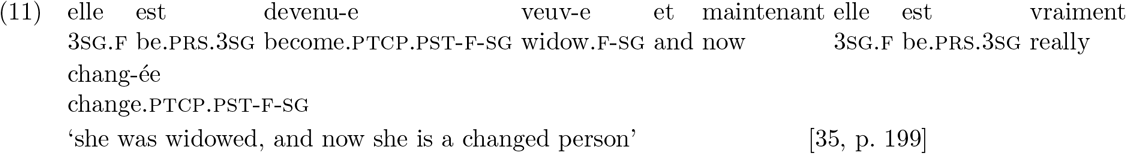

In (10), the coordination construction can convey both a conjunctive and a temporal relation between the two clauses (i.e., after slamming the door, he realized the keys were still inside). The semantics of a construction can be cumulative, allowing multiple layers of meaning to be expressed simultaneously, as in (11), which reflects conjunctive, temporal, and causal interpretations.

The full list of the variables, together with an explanation and a coded construction for example, is shown in Table 1.

**Table 1:**
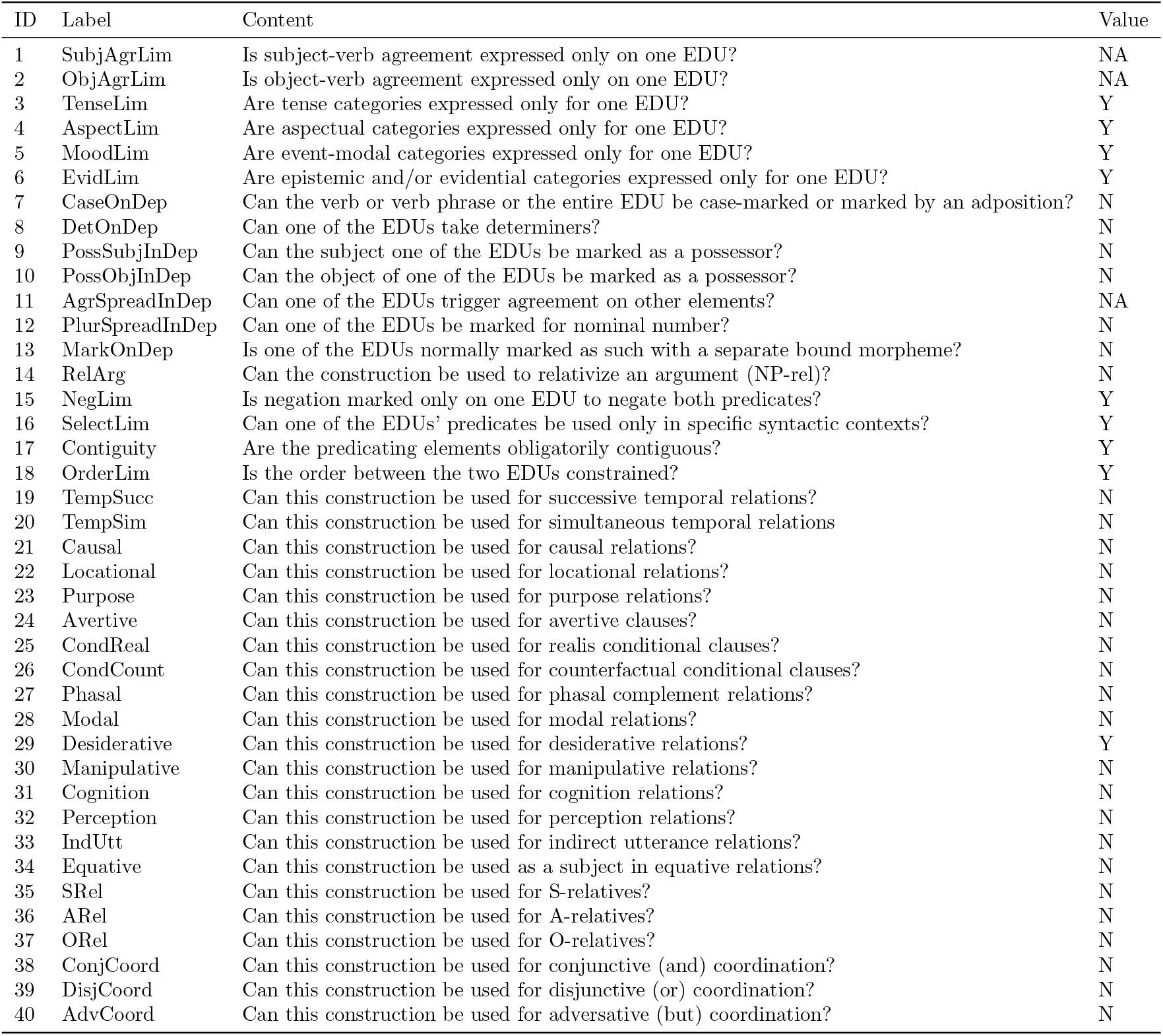
Excerpt from the Burmese (nucl1310) questionnaire used for morphosyntactic and semantic annotation of clause combining constructions. The table shows the coding sheet as it appeared before data transformation for modeling, illustrating construction 19.1, “Desiderative with marker 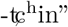 (adapted from Jenny and Hnin Tun [36], esp. pp. 211-212). The variables are coded using Y/N/NA/? values. Variables 1-6 refer to finiteness and de-verbalization properties; variables 7-12 capture noun-like features of event-denoting units (EDUs); variable 13 describes flagging strategies; variable 14 pertains to relativization; variables 15-16 describe structural integration; variables 17-18 address linearization; and variables 19-40 represent semantic relations. For syntactic features (variables 1-18), a “Y” answer indicates a higher degree of syntactic integration.

A straightforward implementation of this coding scheme to study the syntax-semantics interface is to map the semantic relations (IDs 19-40) to those captured in the IRH Van Valin [20]. This mapping is summarized in Table 2, with unmatched categories grayed out.

**Table 2:** Correspondence between the IRH categories and the semantic variables used in this study. Categories in the right-hand column represent the labels used in our dataset for coding constructions. Grayed-out cells indicate relations from the IRH that were not included in our coding scheme.

| ISRH Label | Semantic Label in Dataset |
| --- | --- |
| Causative[1] | Causal |
| Phase | Phasal |
| Manner | Locational |
| Motion | Locational |
| Position | Locational |
| Means | Locational |
| Psych-action |  |
| Purposive | Purpose |
| Jussive |  |
| Causative[2] |  |
| Direct perception | Perception |
| Indirect perception | Perception |
| Propositional attitude |  |
| Cognition | Cognition |
| Indirect discourse | Indirect Utterance |
| Direct discourse |  |
| Circumstances |  |
| Reason |  |
| Conditional | Conditional |
| Concessive |  |
| Simultaneous actions | Temporal Simultaneous |
| Sequential actions | Temporal Successive |
| Situation-situation: unspecified |  |

The correspondence in Table 2 predicts that for any sequential pair of semantic features, syntactic integration should weaken. This is illustrated by simulations in Figure 1, showing how effects would look like if the hierarchy is fully supported. For example, the prediction is that phasal meanings should show less syntactic integration than causal meanings, or succession in time should show less integration that simultenously occuring events. Any contrast above zero, i.e. where a meaning lower on the hierarchy would show more rather than less syntactic integration would violate the predictions. At the same time, the IRH should not be interpreted as overly deterministic, especially given that Van Valin [20, pp. 208–209] acknowledges the complexity of the relationship between syntactic and semantic relations in clause linkage, resulting in deviations from 1:1 mappings. Therefore, the prediction allows for considerable uncertainty and perhaps even some amount of violations.

**Figure 1.**
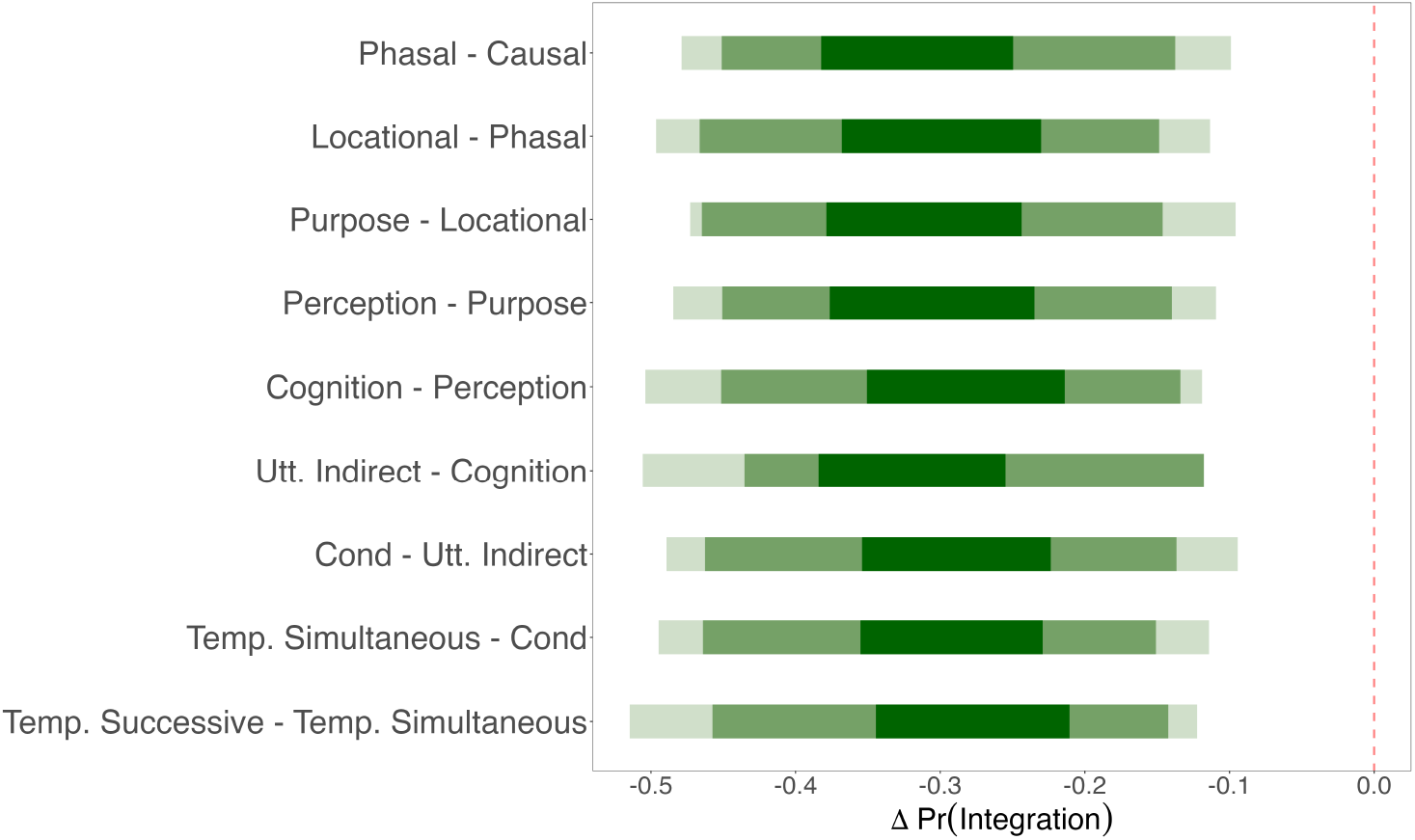
Predicted simulated effects of semantics on syntax in clause linkage. When walking down the hierarchy, each sequential pair of meanings is expected to decrease the probability of integration across the syntactic features of the constructions that express the relevant meanings. The shadings simulate the notion of uncertainty about the estimates (from 95% to 89% to 50% around the median) that is part and parcel of the theory.

Our dataset, which allows testing these predictions, contains 696 constructions drawn from three language families: Indo-European (285 constructions), Sino-Tibetan (300 constructions), and Tupí-Guaraní (111 constructions). The distribution of languages is shown in Figure 2A, and the transformed dataset structure is summarized in Figure 2B. Each row represents a unique combination of semantic and syntactic features within a construction. In addition to these core variables, we include random-effect covariates to account for language relations and areal proximity, thus incorporating phylogenetic and spatial information into our data.

**Figure 2.**
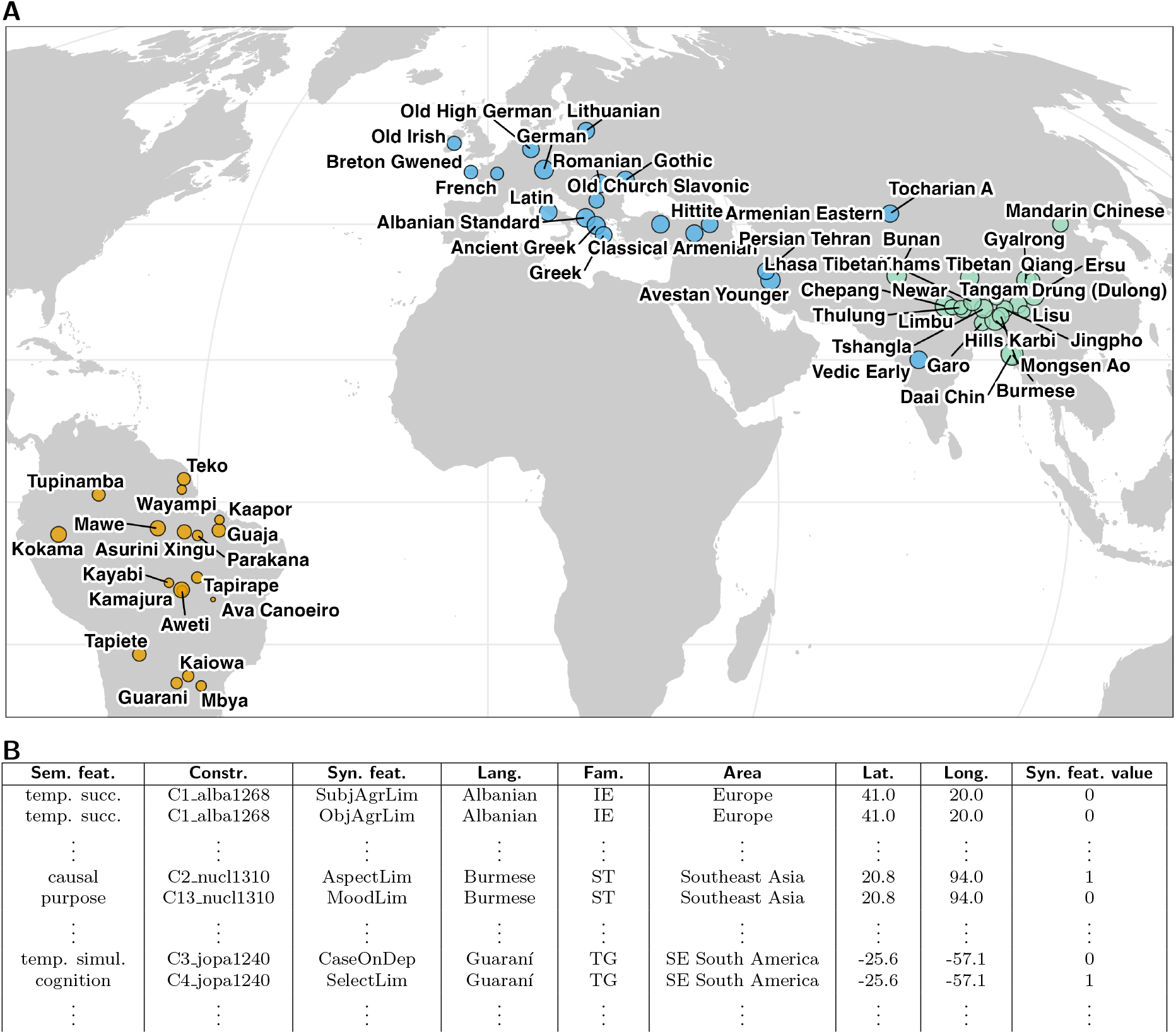
**(A)** Geographic distribution of the languages in our sample, representing three language families: Indo-European (blue), Sino-Tibetan (green), and Tupí-Guaraní (orange). Each circle corresponds to a single language; circle size reflects the number of syntactic constructions known and coded for that language (larger circles represent more constructions). **(B)** Structure of the dataset. Each row represents a unique combination of semantic and syntactic features within a construction. The columns include the semantic value of the construction, a construction ID, the syntactic variable, language name, language family, geographic area (from AUTOTYP), coordinates (from Glottolog) and the syntactic variable value (binary).

## 4 Methods

Our modeling strategy is designed to assess whether a greater degree of semantic integration corresponds to a higher degree of syntactic integration, while taking into account the fact that the languages in our sample have been evolving over time, and may display different propensities toward integration (both in general and at the level of construction type) that are due to their positions in phylogenetic space. Several proposals have been made for addressing phylogenetic structure in linguistic typology. The traditional approach has been to treat phylogeny primarily as a confounding factor — that is, as a historical contingency that obscures rather than informs our understanding of cross-linguistic patterns. From this perspective, diachronic relatedness is viewed as a nuisance variable that should be controlled for rather than examined in its own right.

When treating relatedness as nuisance variable, one strategy is to reduce phylogenetic dependencies already during data collection by sampling languages to be as phylogenetically independent as possible. This facilitates the identification of general cross-linguistic tendencies [see, e.g., 37, 38, 39]. More recently, an alternative approach has gained traction: explicitly incorporating phylogenetic information into statistical models. One way to do this is by modeling family relationships as group-level effects within a hierarchical framework [40, 41]. Still, it remains an open question which level of phylogenetic grouping is most appropriate for analyzing specific linguistic features [42].

We incorporate a variety of hierarchical phylogenetic models into our analyses.^4^ Some of these are in line with the philosophy of treating phylogenetic relatedness as a nuisance variable; elsewhere we build information about predictors of interest directly into the parameters of the evolutionary model in a more “explicit” formulation of the relevant diachronic dynamics [43, 44]. Below, we motivate and describe the approaches we take in detail.

To model the evolution of syntactic features across languages, we first compare two widely used stochastic processes in evolutionary linguistics and biology: Brownian Motion (BM) and the Ornstein-Uhlenbeck (OU) process. BM serves as a baseline model in which traits evolve through an undirected random walk, accumulating changes gradually over time. Under this framework, syntactic divergence between languages is attributed solely to stochastic drift, without any directional forces [45]. Although analytically tractable and conceptually simple, BM lacks mechanisms to account for functional constraints or convergence due to selective pressures, limiting its explanatory power in cases where such forces are hypothesized [46].

Building on the BM framework, the OU process introduces a critical modification: a restoring force that pulls traits toward an optimal value [47]. This makes the OU model particularly suitable for scenarios involving stabilizing selection — cases in which linguistic features are not drifting randomly but are instead biased toward certain preferred syntactic configurations. The dynamics of the OU process are governed by three parameters:

- *α*: the strength of selection, indicating how strongly traits are drawn toward the optimum;
- *θ*: the optimal trait value toward which evolution is biased;
- *σ*: the variance of the stochastic component, reflecting the intensity of random drift.^5^

Conceptually, the OU process generalizes BM: when the strength of selection parameter (*α*) approaches zero, the influence of the restoring force vanishes, and the model reduces to pure Brownian motion [48].

A key advantage of this approach is that it explicitly models the phylogenetic processes. Standard phylogenetic regression models typically treat shared ancestry as a nuisance variable to be statistically adjusted for [49]. In contrast, the OU model treats phylogeny as an integral part of the evolutionary process, incorporating phylogenetic distances directly into the likelihood function [50]. This enables us not only to assess whether related languages resemble one another, but also to investigate the sources of this similarity: whether it results from shared ancestry, convergence toward functional optima, or both.

A further advantage of an explicit model is that it treats present-day syntactic distributions as the outcome of historical evolutionary processes. It does so by leveraging phylogenetic relationships among languages, represented as cophenetic distance matrices derived from posterior samples of phylogenetic trees [51]. These are in turn estimated from lexical cognate replacement rates in a Bayesian framework [52, 53, 54]. For each language family, we fit the model using 50 trees drawn from the posterior distribution, incorporating uncertainty in both branching structure and branch lengths. These trees define a covariance structure among languages based on shared ancestry, which is further shaped by the parameters of the OU model.

To capture variation in evolutionary dynamics, we implement the OU model in a hierarchical structure that allows parameters to vary across syntactic and semantic features. This reflects the assumption that different syntactic constructions are shaped by distinct selective pressures, analoguous to similar differences in biology [48, 55]. For each semantic-syntactic combination, we estimate separate values for the strength of selection (*α*) and the variance of stochastic drift (*σ*), quantifying how strongly traits are drawn toward an optimal configuration and how much they fluctuate due to random processes. These parameters are then used to construct covariance matrices that represent the expected similarity between language pairs as a function of their phylogenetic distance. The matrices are decomposed to generate phylogenetically structured random effects, which, together with additional random intercepts, inform the predicted probability of syntactic integration in each language.

Following the construction of the covariance structure, parameter estimation is conducted within a Bayesian framework using Markov Chain Monte Carlo sampling to approximate the joint posterior distribution over the OU parameters [56]. The likelihood function integrates observed trait values, phylogenetic relationships, and the dynamic covariance matrices derived from the OU model, evaluating how probable the data are under different parameter configurations. The global optimum *θ* is modeled as a hierarchical intercept, with group-specific deviations capturing variation across syntactic and semantic features (see the *Supporting Information*, Section S2, for detailed model specifications).

We fitted separate models for each of the three language families, resulting in a total of three phylogenetically informed regression models. Additionally, we allow the standard deviation by semantic feature to be inferred on the basis of the data, as is standard for random/varying effects. As an alternative, we also fit models where this standard deviation is fixed at 2. This effectively blocks the shrinkage of estimates that arises from treating semantic features as random/varying effects, a phenomenon which could potentially reduce the power of detecting effects of semantics on syntax.

The BM model retains the same hierarchical structure as the OU model, incorporating random/varying effects for language, feature, semantic domain, and construction. It was fitted using the same set of posterior trees to ensure comparability.

Additionally, to probe for potential areal effects, we evaluated three additional configurations of the BM model. We refer to these configurations as “phylogenetic regression” in line with the literature, although they are essentially equivalent to a BM model with a spatial term. One phylogenetic regression model incorporates a Gaussian Process (GP) component to model the effects of areality [42], one uses a spatial random/varying effect [57, 40], and a third one one foregoes any spatial component. Given the complexity of fitting a large number of sensitivity analyses, these models were fitted on the maximum clade credibility tree [58].

The random/varying effect approach was based on the area classification from the AUTOTYP database [59], which groups languages into broad geohistorical regions. This method captures coarse-grained spatial structure by allowing variation to cluster within predefined regional categories. The GP model offers a more fine-grained treatment of geographic influence. Using latitude and longitude coordinates from Glottolog [60], we implemented a bivariate Gaussian Process GP over geographic space. GPs are non-parametric models that capture non-linear relationships among data points by modeling similarity as a function of spatial proximity [61, 62]. In this spatial context, languages located closer together exert more mutual influence than those farther apart. The degree of the spatial smoothing effect is inferred from the data itself, allowing the model to adjust to varying spatial influence across regions. The GP also reflects the uneven density of language samples, yielding narrower uncertainty intervals in well-sampled areas and broader ones where data are sparse. The spatial correlation structure is governed by a covariance function (in our model, the Exponential Quadratic Kernel) which defines how similarity between observations decays with geographic distance. Despite its advantages, the GP approach has a key limitation relative to the area-based model: it implicitly assumes that languages have remained fixed in their geographic locations. In contrast, the area-based model assumes only that languages originate from the same geo-historically (i.e., culturally) defined regions, permitting the possibility of movement within those areas [63].

In addition to models that include phylogenetic and spatial components, we also fitted a baseline model excluding the semantic predictor. This allowed us to assess whether semantics contributes to patterns of syntactic integration at all.

All models, regardless of configuration, included random/varying intercepts for specific constructions to account for idiosyncratic variation across them. Furthermore, we included both random/varying intercepts and random/varying slopes for syntactic features, allowing the effect of semantics to vary across them.

## 5 Results

We conducted model comparisons using leave-one-out cross-validation [64] to evaluate which modeling approach better accounted for the data: random drift (BM) or selection toward an optimum (OU). Across all language families, the best-performing model was the OU model with non-fixed (i.e., estimated) standard deviations, as shown in Tables 3-5. Notably, in every case, models with estimated standard deviations outperformed those with fixed standard deviations.

**Table 3:**
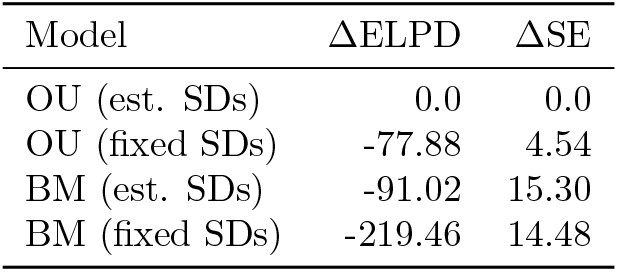
Indo-European.

**Table 4:**
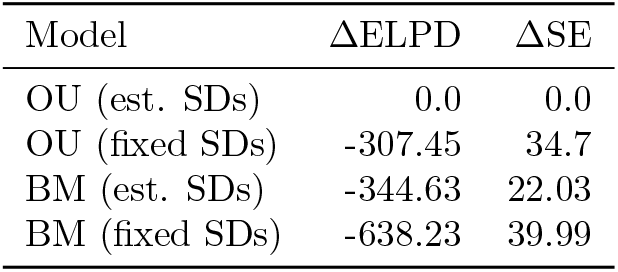
Sino-Tibetan.

**Table 5:**
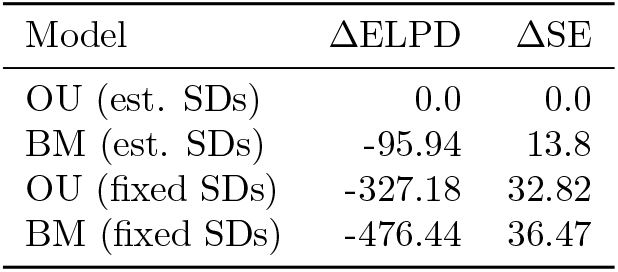
Tupí-Guaraní.

For completeness, we also compared the explicit BM and OU models with the basic phylogenetic regression model spatial effect (Tables 6-8). In all cases, the explicit models outperformed the phylogenetic regression model.^6^ We return to potential effects of spatial locations in Section 5.2 below.

**Table 6:**
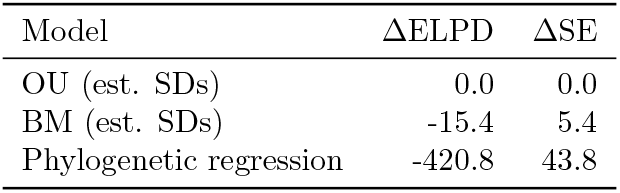
Indo-European.

**Table 7:**
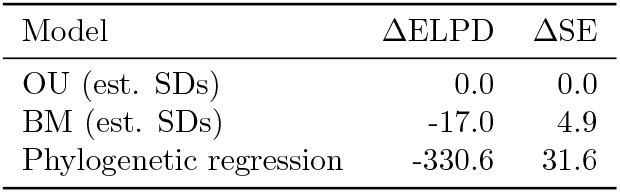
Sino-Tibetan.

**Table 8:**
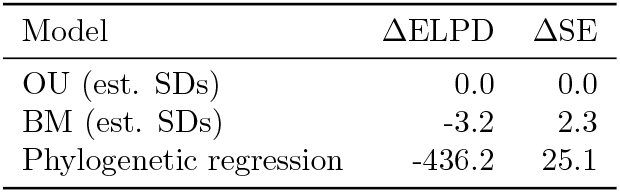
Tupí-Guaraní.

To facilitate interpretation of our model results, we report marginal effects [65], which summarize the influence of predictors while incorporating the full model structure. This approach is especially well-suited for complex models like ours, which include non-linear relationships and hierarchical components. Rather than relying on raw coefficients, which can be difficult to interpret in the presence of interactions or varying slopes, we focus on post-estimation quantities that are more intuitive and substantively meaningful [66, 67]. To assess the effect of the semantics predictor, we computed average marginal contrasts between its levels. Specifically, we generated predicted outcomes from the fitted model for each observation under each level of the semantic relations variable and then calculated the average differences between these predictions. This method integrates over the full distribution of covariates, thereby avoiding misleading interpretations based on fixed or average covariate values [68, 69].

### 5.1 OU models (best fitting)

Figure 3A shows that the predictions of the IRH are rarely supported. This is the case in only three contrasts across all families: Locational meanings show less syntactic integration than phasal meanings in Indo-European; in Sino-Tibetan, complementation under perception verbs shows less syntactic integration than purpose clauses, and successive events are less integrated than simultaneous events. All other contrasts are either non-decisive, with estimates including zero in their 95% highest posterior density interval or go into the opposite direction (e.g. phasal meanings show decisively stronger, not weaker, syntactic integration in Indo-European and Sino-Tibetan).

**Figure 3.**
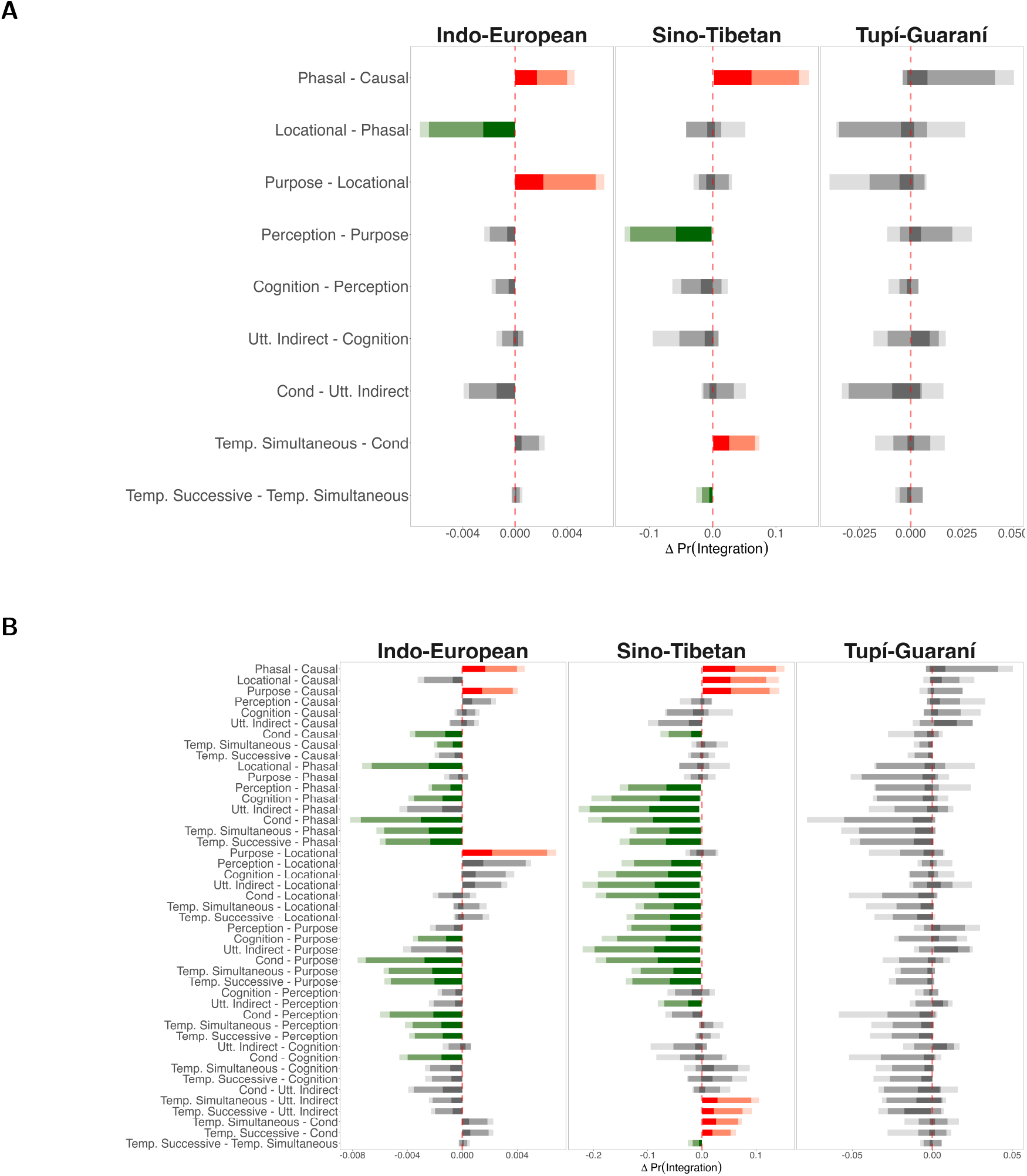
Marginal effects of semantic contrasts estimated from the Ornstein-Uhlenbeck model. Panels show average differences in probability of syntactic integration between semantic relations across constructions, languages, and syntactic features: **(A)** Sequential contrasts following the IRH in a strict sense; **(B)** Pairwise contrasts allowing effects between nonadjacent meanings in the IRH. The intervals represent the 95%, 89%, and 50% highest probability density. Contrasts decisively supporting the IRH are colored green; those violating it are in red. We assume evidence is decisive if 0 is outside the 95% highest posterior density interval.

Substantial variation emerges across language families. In Indo-European, three of the nine sequential contrasts have decisive sizes, while in Sino-Tibetan, four of nine have decisive sizes. In contrast, none of the sequential comparisons in Tupí-Guaraní are decisive, suggesting a weaker or more diffuse evolutionary signal in this family.

In addition to the sequential contrasts based on the predictions of the IRH, we conducted a full set of pairwise comparisons across the semantic hierarchy (Figure 3B). This loosens the interpretation of the IRH as a strict sequence, allowing for the possibility of meanings that are non-adjacent on the hierarchy to differ in syntactic integration while effects in between show no decisive difference. In Tupí-Guaraní, these comparisons revealed no decisive differences, consistent with the sequential results. In contrast, Indo-European and Sino-Tibetan showed numerous additional decisive contrasts, but only some are consistent with the predictions of the IRH, others contradicting them. Specifically, 19 out of 45 contrasts were decisive in Indo-European (3 against the IRH), and 28 in Sino-Tibetan (7 against the IRH).

### 5.2 Phylogenetic regression

For completeness, we also report the results of the phylogenetic regression model; though it showed the worst model performance, this approach is increasingly common in typology, and it is useful to evaluate the role of spatial structure and contact-induced change in shaping the syntax-semantics interface Tables 9-11.

**Table 9:** Indo-European.

| Model | ΔELPD | ΔSE |
| --- | --- | --- |
| Spatial random effect | 0.0 | 0.0 |
| Spatial Gaussian process | -0.59 | 0.31 |
| No spatial effect | -0.86 | 0.30 |
| No semantic effect | -1163.36 | 43.94 |

**Table 10:** Sino-Tibetan.

| Model | ΔELPD | ΔSE |
| --- | --- | --- |
| Spatial Gaussian process | 0.0 | 0.0 |
| No spatial effect | -1.47 | 0.94 |
| Spatial random effect | -1.74 | 0.93 |
| No semantic effect | -1067.48 | 38.01 |

**Table 11:** Tupí-Guaraní.

| Model | ΔELPD | ΔSE |
| --- | --- | --- |
| No spatial effect | 0.0 | 0.0 |
| Spatial random effect | -0.14 | 0.24 |
| Spatial Gaussian process | -0.15 | 0.30 |
| No semantic effect | -269.12 | 22.67 |

In Indo-European, the model with spatial random/varying effects performed best, though its advantage over the Gaussian process model was weak. In Sino-Tibetan, the Gaussian process model slightly outperformed the alternatives, but again, the differences are minor. For Tupí-Guaraní, the model without any spatial component provided the best fit, though differences with the spatial models were minimal. In all families, models including semantic predictors substantially outperformed those without them. At the same time, at the level of individual meaning relations, there are even fewer decisive effects of semantics on syntax than in the OU model (Figure 4). Only the difference between complementation with perception verbs and purposes clauses in Sino-Tibetan remains in line with the IRH predictions. All other contrasts fail to support the IRH predictions.

**Figure 4.**
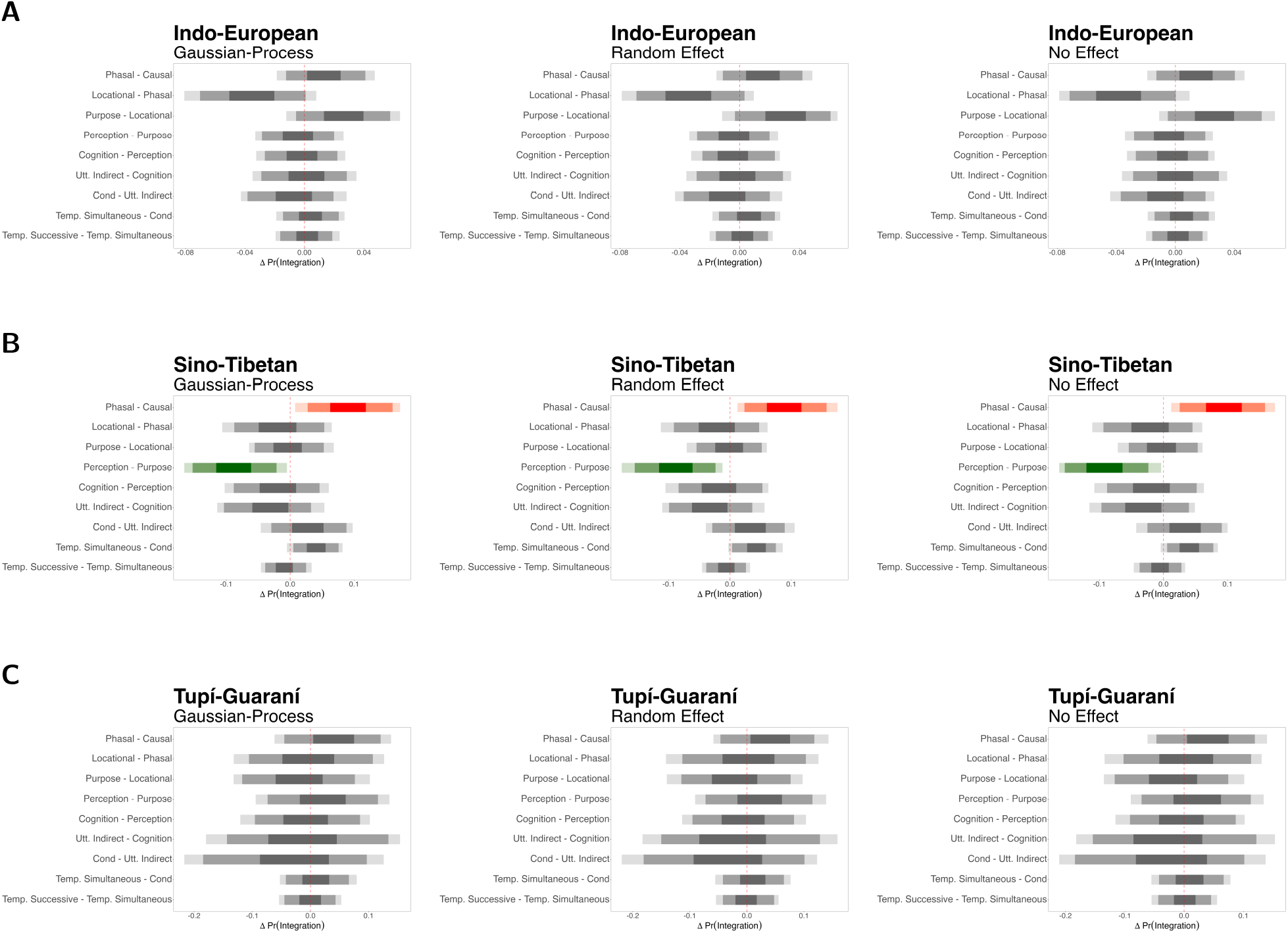
Estimated marginal effects of the semantic predictor across language families and spatial model specifications. Each panel corresponds to a language family: **(A)** Indo-European, **(B)** Sino-Tibetan, and **(C)** Tupí-Guaraní. Within each panel, the columns display results from models controlling for spatial structure in different ways: using a Gaussian Process (left), a spatial random/varying effect by AUTOTYP area (center), and no spatial component (right). The plots show average comparisons between levels of the semantic predictor, with intervals representing posterior uncertainty (95%, 89% and 50% highest posterior density intervals). Across all three families, the choice of spatial modeling has minimal impact on the estimated effect of semantics, suggesting that geographic factors do not substantially influence the relationship between semantic category and syntactic integration. Plotting conventions like in Figure 3.

The choice of spatial modeling has minimal influence on the estimated effect of semantics. Geographic factors do not appear to meaningfully alter the relationship between semantics and syntactic integration.

As noted, while effective in accounting for phylogenetic non-independence, the phylogenetic regression approach does not capture the underlying dynamics of language change. It identifies whether a phylogenetic signal exists but remains agnostic about the direction, rate, or nature of diachronic processes. The largely null results of our regression analyses suggest that the signal of interest may reside not in static phylogenetic relations, but in the temporal dynamics of linguistic evolution itself — an interpretation supported by the OU analysis.

## 6 Discussion

According to the predictions of the IRH, the contrast distributions shown in both panels of Figure 3 should have been consistently negative, as the contrasts were always calculated by subtracting the semantic level higher in the IRH from the one lower. Even allowing for some violations and considerable uncertainty, the complete absence of decisive contrasts in many parts of hierarchy, and for all parts in Tupí-Guaraní, is unexpected.

Some of the decisive contrasts we found even run counter to the predictions of the IRH, particularly those involving the causal semantic level. Van Valin [20] notes that the semantic relations at the upper end of the hierarchy are often lexicalized and not expressed through complex syntactic constructions. As a result, the tightest syntactic linkages may not correspond to the highest-level semantic relations. Even so, our sample includes both lexicalized constructions (e.g., Hittite predicates with *-nu-* or Mandarin serial verb constructions) and non-fully lexicalized causal constructions (e.g., Latin *ut* + subjunctive clauses and Modern Greek *kano* + subjunctive constructions). Yet, in both cases, the observed contrasts still fail to align with the predictions of the IRH. Furthermore, several contrasts that contradict the IRH are also found toward the lower end of the hierarchy, particularly those involving temporal semantics.

The fact that some of these contrasts are limited to specific language families, combined with the presence of several patterns that diverge from the predictions of the IRH, calls into question the universality of the proposed semantic hierarchy. These findings indicate that the syntax-semantics interface may have evolved along lineage-specific paths shaped by historical contingencies, such as schismogenesis and other sociocultural factors [70, 71]. Nevertheless, the observation that most decisive contrasts are shared between Indo-European and Sino-Tibetan suggests a possible, albeit weak, universal tendency to align semantic and syntactic integration — even if not in the manner predicted by the IRH. More research is needed to unravel potentially local trends in this area.

An additional finding concerns the role of geography. Our analysis reveals little to no effect of geographic proximity on the syntax-semantics interface in clause linkage, suggesting that the relevant structural properties are largely insulated from areal diffusion. This conclusion holds across multiple spatial modeling strategies: none of the tested approaches significantly altered the predictions obtained from marginal effect estimates, indicating a general absence of geographic structuring in the data. The broader theoretical implication of these findings is that, while contact-induced change is undoubtedly a significant factor in language evolution and typology, and thus must be accounted for in cross-linguistic analyses, its impact may be highly dependent on the linguistic domain in question. At least for properties situated at the syntax-semantics interface, as operationalized in this study, contact appears to be a secondary force compared to language-internal, system-driven processes. This finding suggests that the syntax-semantics interface is particularly resistant to borrowing and contact-induced change. Prior research has similarly argued that properties at this interface are less susceptible to diffusion across language boundaries, due to their deep embedding in grammatical architecture, their close ties to language-internal constraints, and their lack of easily-learnable over signals in morphology or phonology [72, 73]. Our results lend empirical support to this claim, reinforcing the idea that syntactic integration reflects more language-specific grammatical systems rather than areal influences. This resistance to borrowing underscores the deeply rooted nature of interface phenomena and supports the broader view that not all grammatical domains are equally permeable to areal influence. Of course, more languages and families need to be tested before any firm conclusions. We cannot exclude the possibility that areal effects re-emerge at either a global scale or in specific regions of the world.

A secondary, methodological point arises when contrasting the outcomes between the two modeling approaches. The phylogenetic regression yielded minimal effects, while the OU model revealed stronger and more consistent patterns. The key difference lies in how each model treats phylogeny. Phylogenetic regression incorporates relatedness as a background covariance structure, controlling for shared ancestry but not explicitly modeling evolutionary processes. This can obscure meaningful signals by treating phylogeny as statistical noise rather than as a mechanism of trait evolution. As a result, the weak effects observed may reflect the model’s limited capacity to detect adaptive variation, rather than a true absence of signal. This highlights the importance of modeling phylogeny explicitly — that is, with approaches that incorporate branch lengths, allow for heterogeneous evolutionary rates, and estimate the full joint likelihood of the data.

## 7 Conclusions

This study provides evidence for only a weak relationship between semantic properties and syntactic integration in clause linkage. The strength and nature of these effects vary significantly across language families, suggesting that semantic-syntactic mappings are mediated by lineage-specific evolutionary trajectories. This highlights the importance of considering phylogenetic context when assessing cross-linguistic generalizations.

At the same time, we find no evidence for a major impact of language contact on semantic-syntactic mapping. This is consistent with the notion that contact primarily affects patterns that are readily signaled by overt markers and does not impact the syntax/semantics interface, which is more tightly integrated with the internal mechanics of grammar.

Methodologically, our study demonstrates the critical importance of fully explicit phylogenetic modeling. Traditional approaches, which often treat phylogeny as a background or nuisance variable, risk obscuring meaningful patterns. In contrast, dynamic evolutionary models such as the OU framework enable researchers to model trait evolution directly, accounting for both historical signal and structural constraint. Our use of such models revealed patterns that were not detectable using standard phylogenetic regression techniques. This underscores the value of integrating evolutionary theory more deeply into the toolkit of linguistic typology and calls for a reassessment of established methodological practices in the field.

Despite these contributions, several limitations must be acknowledged. Our sample includes only three major language families, Indo-European, Sino-Tibetan, and Tupí-Guaraní, and since our results reveal distinct evolutionary trajectories for each, incorporating additional families is essential to capture a broader spectrum of evolutionary dynamics. Furthermore, although our feature coding is based on established typological criteria, some semantic distinctions may not be fully represented in all language-specific descriptions, potentially introducing bias.

Taken together, our findings point to a more nuanced understanding of the syntax-semantics interface. Rather than supporting rigid hierarchies, the evidence favors a model in which cross-linguistic patterns emerge from the interaction of broad functional pressures and lineage-specific grammatical developments. This perspective encourages a shift away from universalist assumptions toward models that accommodate both cross-linguistic tendencies and historical contingency.

## Acknowledgements

We are grateful to Mathias Jenny (Sino-Tibetan), Rik van Gijn (Tupí-Guaraní), and Paul Widmer (Indo-European) for their help with data collection.

## Contributions

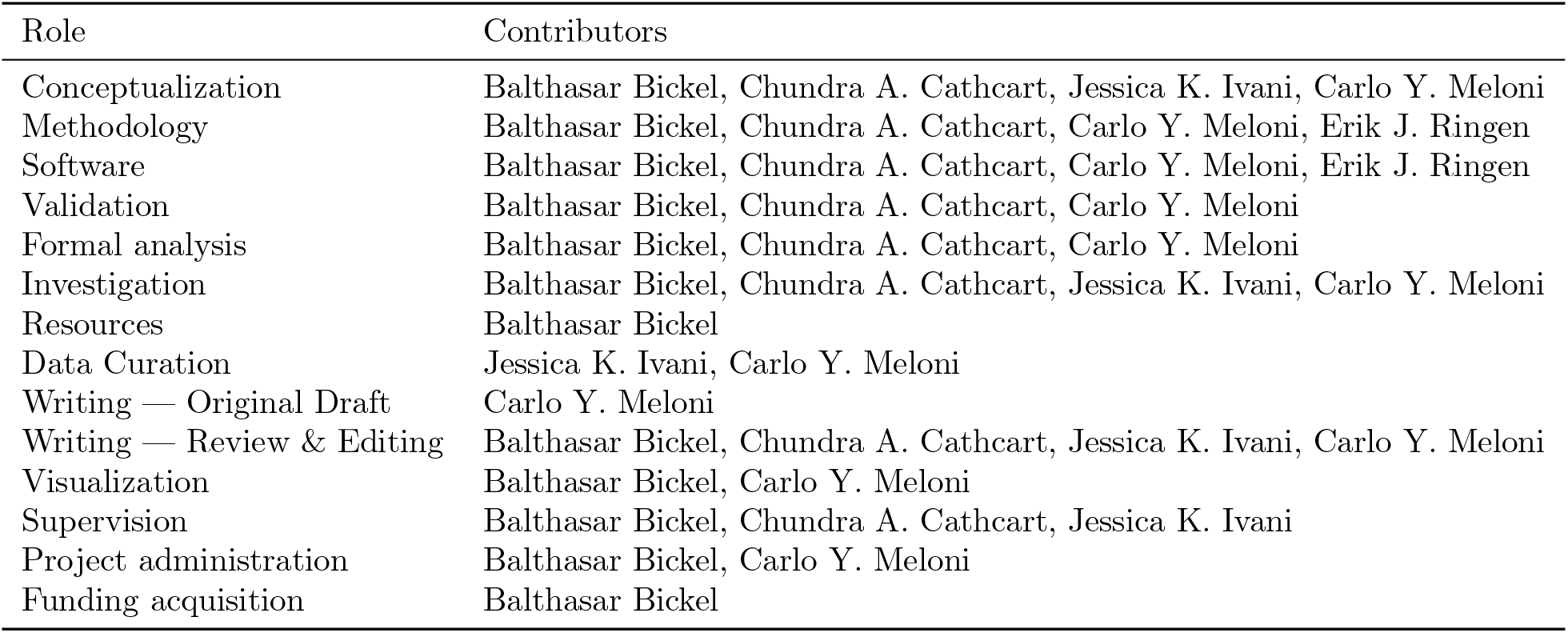

## Supplementary Information

## S1 Phylogenetic Regression

We fitted a Bayesian phylogenetic Bernoulli regression model using the brms interface [74] to Stan [75] in R [76]. The model was designed to examine the relationship between the semantics of a clause linkage construction and the presence of (morpho)syntactic constraints, while accounting for both phylogenetic non-independence and random variation across syntactic constructions and syntactic features.

### S1.1 Data and Preprocessing

The data consists of binary observations *y*_*i*_ ∈ {0, 1}, where each *y*_*i*_ denotes whether a given (morpho)syntactic constraint is present in a language. Each data point is associated with:

- **Semantic level** (sem_*i*_): a categorical predictor encoding the semantic relation expressed by the construction.
- **Syntactic feature** (feat_*i*_): a categorical variable identifying the syntactic constraint being analyzed.
- **Syntactic construction ID** (cons_*i*_): a grouping variable indexing the construction with the feature and semantics.
- **Language identifier** (phylo_*i*_): a grouping variable matching the language to the tips of a phylogenetic tree.

The model was fitted separately for Indo-European, Sino-Tibetan, and Tupí-Guaraní. For each family, we used a Maximum Clade Credibility (MCC) tree derived from posterior tree samples [52, 54, 53]. The MCC trees were converted into variance-covariance matrices using the function vcv.phylo() from the ape package [77].

### S1.2 Mathematical Formulation

The model assumes:

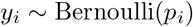

With

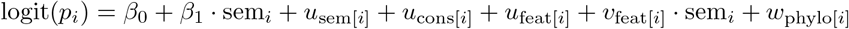

Where:

- *β*_0_ *∼ N* (0, 2) is the population-level intercept. *N*
- *β*_1_ *∼ N* (0, 2^2^) is the fixed effect of the semantic level.
- 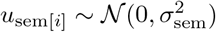 is a random intercept for semantic level.
- 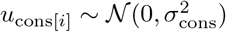is a random intercept for constructions.
- 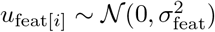 is a random intercept for syntactic features.
- 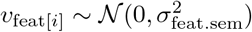 is a random slope of semantic level per feature.
- 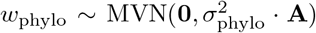 is a phylogenetic random effect, where **A** is the phylogenetic variance-covariance matrix derived from the MCC tree.

This formulation corresponds to the following brms model syntax:

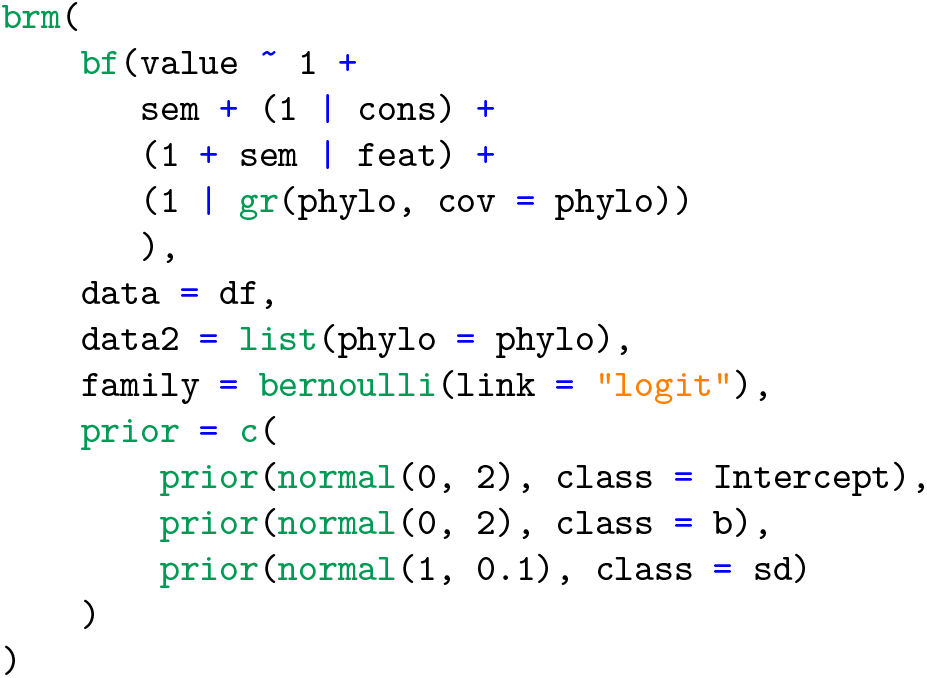

### S1.3 Phylogenetic Term and MCC Tree

The phylogenetic effect was implemented using the `gr()` function in `brms`, which allows for random effects to follow a known covariance structure:

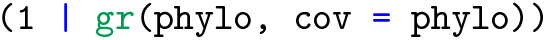

This formulation introduces a multivariate normal prior over language-level intercepts, with a covariance structure proportional to the cophenetic distance matrix **A**. The matrix **A** captures the historical signal shared among languages based on their evolutionary relationships. Closely related languages are expected to exhibit more similar intercepts than distantly related ones.

Our approach extends that of Guzmán Naranjo and Becker [42], who construct a similarity matrix using taxonomic trees from Glottolog [60] and define micro-family groupings. We use phylogenetic, not taxonomic trees, and allow similarities to vary with the branch lengths estimated in published phylogenies on the basis of cognacy replacement rates and extralinguistic calibration points [52, 54, 53].

### S1.4 Controlling for Spatial Autocorrelation

To control for potential geographic confounds, we implemented two additional models incorporating spatial effects:

#### Model with Micro-Area Random Effects

This model includes random intercepts over the geohistorically defined areas from AUTOTYP [59]. It accounts for areal clustering by allowing local spatial dependencies:

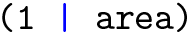

#### Model with a Gaussian Process over Geography

The second spatial model incorporates a continuous spatial effect using a Gaussian Process (GP) over the geographic coordinates (longitude and latitude) of each language [taken from 60]:

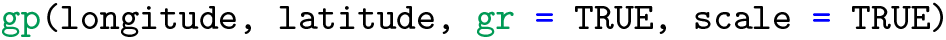

This approach captures smooth spatial variation across geographic space, without requiring discretization into areas. The GP estimates a covariance function over spatial distances, allowing the model to capture spatial autocorrelation that may reflect contact zones or diffusion. This implementation follows the method described in Guzmán Naranjo and Becker [42, pp. 630–633], where spatial structure is modeled using a squared exponential kernel function. The GP term introduces a distance-based correlation across languages, with the strength of correlation decaying as a function of geographic separation. This enables the model to capture areal effects that are not explained by phylogenetic relatedness.

## S2 Hierarchical Brownian Motion Model

Additionally to the brms-based phylogenetic regression, we implemented a Bayesian binary logistic regression model directly in Stan [75], which explicitly specifies the internal structure of the phylogenetic effect via a Brownian motion (BM) GP. While the two models are conceptually similar in treating languages as non-independent samples, the model described in this section fully specifies in the internal structure of the phylogenetic component.

### S2.1 Mathematical Formulation

Each binary observation *y*_*i*_ ∈ {0, 1} is modeled as *y*_*i*_ *∼* Bernoulli(*p*_*i*_), with logit(*p*_*i*_) = *ψ*_*i*_.

The linear predictor *ψ*_*i*_ is composed of several additive components:

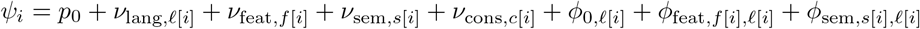

Where:

- *p*_0_ *∼ N* (0, 1) is global intercept.
- 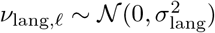is the language-level random effects.
- 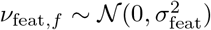 is syntactic feature-level random effects.
- 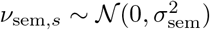 is the semantic-level random effects.
- 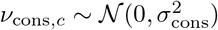 is the construction-level random effects.

All random effects are non-centered: they are defined as standard normal variables scaled by their respective standard deviations to improve sampling efficiency in hierarchical settings [78, p. 319].

### S2.2 Phylogenetic Term

Three types of phylogenetic effects are included:

- A global BM effect 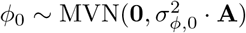, shared across all observations.
- Feature-specific BM effects 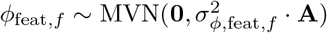, one for each feature.
- Semantic-specific BM effects 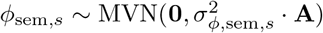, one for each semantic group.

Each covariance matrix **A** is a patristic distance matrix derived from a phylogenetic tree. To account for uncertainty in the phylogeny, the model was fitted both to a single MCC tree (used for comparison with the phylogenetic regression), and to a sample of 50 trees, allowing the phylogenetic effect to be marginalized across multiple plausible evolutionary histories.

For the feature- and semantics-specific effects, evolutionary rate variation is modeled hierarchically:

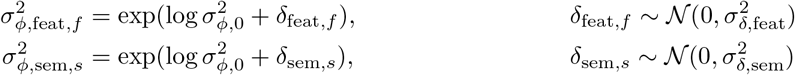

This formulation of the phylogenetic component allows for lineage-specific, feature-specific, and semantic-specific rates of diffusion along the phylogeny, enabling the model to capture heterogeneous patterns of historical signal in the data.

The BM model described here did not contain a spatial component, since its inclusion did not affect the predictive performance in what we call the phylogenetic regression model (Section S1).

## S3 Hierarchical Ornstein–Uhlenbeck Model

To complement the models described above, we also implemented a regression model with a fully specified Ornstein–Uhlenbeck (OU) process for the phylogenetic component. The OU process extends the BM framework by incorporating stabilizing selection toward an optimal trait value, allowing us to model phylogenetic signal that decays with evolutionary distance.

### S3.1 Mathematical Formulation

As in the BM models, each observation *y*_*i*_ ∈ {0, 1} is assumed to follow a Bernoulli distribution with success probability *p*_*i*_, linked to a linear predictor *ψ*_*i*_ via the logit function:

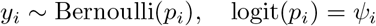

The predictor *ψ*_*i*_ includes fixed and random components, along with three phylogenetic terms:

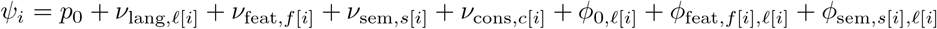

The interpretation and specification of *p*_0_, *ν*_*•*_, and the use of non-centered parameterization follow the structure described in the BM model.

### S3.2 Ornstein–Uhlenbeck Phylogenetic Process

The key difference in this model lies in how the phylogenetic effects *ϕ* are modeled. Instead of assuming that traits evolve by random drift (as in BM), we use a stationary OU process, which introduces a tendency for traits to revert toward a long-run optimum (the global intercept *p*_0_).

The OU process is defined by the stochastic differential equation:

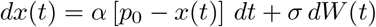

where

- *α* governs the strength of attraction to the optimum *p*_0_,
- *σ* controls the magnitude of random fluctuations,
- *W* (*t*) is a standard Wiener process.

In its stationary form, the OU process induces a GP over languages with the following covariance function [67, p. 482]:

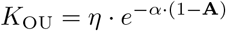

where 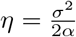 represents the stationary variance of the process.

This formulation ensures that closely related languages share more of their trait values, but that this similarity decays exponentially with evolutionary distance, governed by *α*. In contrast to BM, which implies infinite diffusion over time, the OU model limits long-term divergence by pulling traits toward a central value.

### S3.3 Hierarchical OU Parameters

Each of the three OU components in the model is given its own covariance structure based on this kernel. The parameters *η* and *α* for each feature and semantic group are modeled hierarchically:

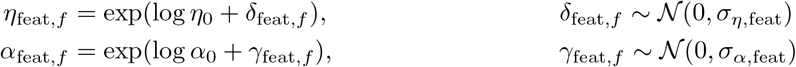

with analogous formulations for the semantic-level parameters *η*_sem,*s*_, *α*_sem,*s*_. The global OU process uses the shared hyperparameters *η*_0_ and *α*_0_, directly applied to the phylogenetic covariance.

### S3.4 Relationship to Brownian Motion

The OU process generalizes BM. As *α →* 0, the OU kernel converges to the BM kernel:

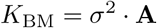

This means that the OU model can flexibly capture both drift-like and constraint-like patterns of trait evolution, depending on the inferred value of *α*. When the phylogenetic signal is weak or traits are highly conserved, higher values of *α* shrink the phylogenetic contribution toward zero.

## Footnotes

1 We do not claim exhaustivity. See also Silverstein [4], Haiman [5, 6, 7], Haiman and Thompson [8], and Newmeyer [9] for discussion on iconicity in clause linkage and syntax in general; Bolinger [10] for the role of intonation in clause combining, especially subordination; Hengeveld [11] on adverbial clauses; Noonan [12] on complementation.

2 Numerous counterexamples have been discussed in the grammaticalization literature: see Mithun [15] for perception-cognition-utterance verbs grammaticalized as evidentials; and see also Heine, Claudi, and Hünnemeyer [16], Traugott and Heine [17], and Givón [18] especially with respect to complementation and co-lexicalization.

3 Nine types are the result of the combination between nexus and juncture types. Additionally, Van Valin [22] introduces sentential coordination and sentential subordination.

4 The codes for the models, along with the data and additional scripts, are provided in the Zenodo repository.

5 Our implementation of the model captures the evolutionary processes through a phylogenetic covariance matrix where *σ* is scaled by *α*, representing the stationary (long-term) variance (*η*) of the process. See below and Supporting Information.

6 Note that the ΔELPD values between the BM and OU models in Tables 3–5 differ from those reported in Tables 6–8.

This discrepancy arises because the models reported in Tables 6–8 were fitted to a maximum clade credibility tree for direct comparability with the phylogenetic regression model, whereas the models in Tables 3-5 were fitted to a sample of trees.

